# Influence of HLA class II polymorphism on predicted cellular immunity against SARS-CoV-2 at the population and individual level

**DOI:** 10.1101/2020.12.24.424326

**Authors:** Hannah C Copley, Loren Gragert, Andrew R Leach, Vasilis Kosmoliaptsis

## Abstract

Development of effective adaptive immune responses after coronavirus disease 2019 (COVID-19) and after vaccination against SARS-CoV-2 is predicated on recognition of viral peptides, presented in the context of HLA class II molecules, by CD4^+^ T-cells. We capitalised on extensive high resolution HLA data deposited in the National Marrow Donor Program registry to obtain detailed information on human HLA haplotype frequencies of twenty five human populations and used a bioinformatics approach to investigate the role of HLA polymorphism on SARS-CoV-2 immunogenicity at the population and at the individual level. Within populations, we identify wide inter-individual variability in predicted CD4^+^ T-cell reactivity against structural, non-structural and accessory SARS-CoV-2 proteins, according to expressed HLA genotype. However, we find similar potential for anti-SARS-CoV-2 cellular immunity at the population level, across all ethnic groups examined, suggesting that HLA polymorphism is unlikely to account for observed disparities in clinical outcomes after COVID-19 among different race and ethnic groups. We predict robust immune reactivity against SARS-CoV-2 Spike protein, the basis for the majority of current vaccination efforts, both at the population and individual level, despite significant variation in Spike-derived peptide presentation by individual HLA genotypes. Finally, we provide comprehensive maps of SARS-CoV-2 proteome immunogenicity accounting for population coverage in major ethnic groups. Our findings provide important insight on the potential role of HLA polymorphism on development of protective immunity after SARS-CoV-2 infection and after vaccination and a firm basis for further experimental studies in this field.

## Introduction

The new severe acute respiratory syndrome coronavirus 2 (SARS-CoV-2), responsible for coronavirus disease 2019 (COVID-19), has caused an ongoing pandemic with 75,704,857 confirmed cases and 1,690,061 deaths worldwide (as of 21 December 2020^1^). Several risk factors for severe COVID-19 are now well-established, including age, gender, obesity and various co-morbidities such as diabetes, cancer and cardiovascular or chronic lung disease^2, 3, 4, 5^. There is, however, an urgent need to better understand the role of race and ethnic differences on health outcomes. Several studies have highlighted a disproportionate prevalence of COVID-19 infections, higher rates of hospitalisation, and increased incidence of death in people from black and minority ethnic groups but the underlying reasons for these observations are not well understood^2, 6, 7, 3, 9^.

Even after accounting for well-known risk factors, there is still wide inter-individual clinical variability of COVID-19 outcomes within considered risk groups, which may reflect underlying genetic differences^10^. The principal genetic region involved in immunity against viral pathogens is the Major Histocompatibility Complex encompassing the Human Leukocyte Antigen (HLA) loci. HLA class I (HLA-A, -B, -C) and HLA class II (HLA-DR, -DQ, -DP) proteins present viral peptides for recognition by CD8^+^ and CD4^+^ T-cells, respectively. The latter orchestrate adaptive anti-viral immunity and drive B-cell activation and maturation for robust humoral responses. Extensive polymorphism is observed in the HLA system, resulting in differences in HLA allele frequency both within and across human populations. HLA genotype can be a determining factor in development of protective immunity and, in turn, may account for part of the observed heterogeneity in measured immune responses and in clinical outcomes after SARS-CoV-2 infection^11, 12, 13^. It is also well established that there is marked biological variation in how individuals respond and maintain immunity after vaccination which is, in part, attributable to genetic factors^14^. It is, therefore, important to consider the role of HLA polymorphism when designing viral subunit or peptide vaccine formulations and in assessing population coverage and likelihood of immune protection after vaccination^15^.

When considering the role of the HLA system in COVID-19 susceptibility and vaccine responses, it is essential to account for differences in HLA allele frequencies across human populations and, importantly, for the linkage disequilibrium between HLA loci that result in population-specific haplotype frequencies. Here, we utilise information on human HLA haplotype frequencies of twenty five human populations (four broad population categories and twenty one detailed population subcategories) at an unprecedented scale, capitalising on the extensive high-resolution HLA data deposited in the National Marrow Donor Program Registry, to compute population level immune responses against SARS-CoV-2 based on predicted high-affinity binding of viral proteome derived peptides by HLA class II molecules. Overall, we find similar potential for anti-SARS-CoV-2 cellular immunity across all populations examined suggesting that HLA polymorphism is unlikely to account for observed disparities in clinical outcomes after COVID-19 among different race and ethnic groups. However, within populations, we identify wide variability among individuals in predicted CD4^+^ T-cell reactivity against structural, non-structural, and accessory SARS-CoV-2 proteins, according to HLA genotype. Nevertheless, we predict robust immune reactivity against the SARS-CoV-2 Spike protein, the basis for the majority of current vaccination efforts, both at the population and at the individual level.

## Methods

### SARS-CoV-2 viral proteome and identification of potential T-cell epitopes

Full viral proteome sequences for SARS-CoV-2 were downloaded from UniProt^16^. FASTA-formatted protein sequence data for each protein and protein class were examined individually and in combination. We produced potential peptides of 15 amino acids length (15mers), using sliding windows over the entire proteome. Proteins of fewer than 15 amino acids in length were not examined. Analyses were performed considering proteins individually, some protein domains individually, and in groupings of proteins (both the whole proteome, and all structural, non-structural, and accessory proteins).

### HLA population frequency computation

HLA population frequencies were obtained from US unrelated stem cell donor registry National Marrow Donor Program (NMDP) / Be The Match. High resolution HLA Class II haplotype frequencies (DRB1, DRB3/4/5, DQA1, DQB1, DPA1 and DPB1 loci) were estimated using an expectation-maximization algorithm (as described by Gragert et al. 2013^17^) utilising a cohort of 8.9 million US volunteer donors (NMDP/Be The Match registry snapshot 29/05/2020) HLA typed by molecular methods (supplementary Table 1 showing number of individuals genotyped). US population categories were developed based on a race/ethnicity questionnaire included on the donor consent form. There were four broad population categories (European American, African American, Asian or Pacific Islander and Hispanic) and 21 detailed populations subcategories (African American, African Black, South Asian Indian, American Indian - South or Central American, Alaska native of Aleut, North American Indian, Caribbean Black, Caribbean Hispanic, Caribbean Indian, European Caucasian, Filipino, Hawaiian or other Pacific Islander, Japanese, Korean, Middle Eastern or North Coast of Africa, Mexican or Chicano, Chinese, Hispanic - South or Central American, Black - South or Central American, Southeast Asian, Vietnamese). Some individuals had membership only within a broad population category.

Following Hardy-Weinberg equilibrium proportions, multi-locus HLA Class II genotypes were generated by randomly sampling two haplotypes from the same population HLA haplotype frequency distribution. Simulated genotypes were generated for each of the four broad and 21 detailed population groups, with ten replicates at each population of size 1,000, 5,000 and 10,000.

For population-based analyses at the haplotype level, we analysed haplotypes up to 99% cumulative coverage within each population. HLA alleles, when examined individually, included all HLA alleles present in any of these selected haplotypes for the four broad and 21 detailed population groups.

### Computational peptide-HLA prediction model for T-cell epitope selection

Peptide binding affinity was assessed for all HLA that featured in haplotypes in any of the ethnic populations studied using NetMHCpan v4.0^18^. Peptides were examined both for their predicted binding affinity (nM) and their percentage rank (compared to a pool of representative peptides for that HLA). Peptides were counted as binding if their binding affinity was equal to or less than 500nM and their percentage rank was equal to or greater than 2% (default parameter for strong HLA class II peptide binders). The NetMHCIIpan-4.0 programme also identifies the predicted 9-mer binding core. Total predicted peptide counts for an individual HLA per protein or per whole proteome were calculated by counting peptides only if they were both unique to one another (i.e a unique 15-mer), but also if the predicted binding core (9-mer) was also unique. This prevented the count from appearing falsely elevated due to sequential overlapping peptides which were presenting the same core, from being counted twice. Total predicted peptide counts for an HLA haplotype or genotype similarly required the peptide and the peptide binding core to be unique, except in this case the peptide could be presented in any of the HLA expressed in the haplotype or genotype.

### Statistical analysis

Classification performance of peptide-MHC scoring models was calculated using scikit-learn^19^ in Python using the sklearn.metrics.roc_auc_score (AUROC), sklearn.metrics.average_precision_score (Average Precision), sklearn.metrics.accuracy_score (Accuracy), and sklearn.metrics.classification_report (Sensitivity and Specificity) functions. Population comparisons of peptide scores were performed by calculating the mean and standard deviation using NumPy in Python, and between repeat population simulations using statistics.shaipro (Shapiro-Wilk test for normality) and statistics.kruskal (Kruskal–Wallis one-way analysis of variance) both in Python.

## Results

### Experimental approach

The principal objective of our study was to examine whether genetic variation at the HLA loci may influence immune responses, and therefore COVID-19 clinical outcomes or response to vaccination, among patients from different ethnic populations. HLA class II molecules present peptides from exogenous antigens for CD4^+^ T-cell recognition and are therefore critical components for an effective adaptive immune response that incorporates humoral (B-cell) and cytotoxic (CD8^+^ T-cell) arms.

We analysed peptide binding for all classical HLA class II loci (HLA-DRB1, -DRB3/4/5, -DQA1, -DQB1, and -DPA1, -DPB1). The main analysis focused on four broad population categories, African Americans (AFA), European Caucasians (CAU), Hispanics (HIS) and Asian/Pacific Islanders (API) and sub-analysis included 21 detailed population subgroups of these broad categories. To account for linkage disequilibrium between HLA class II loci, we analysed HLA haplotypes rather than assuming allele frequencies were independent at each locus. HLA haplotype frequencies were estimated utilising HLA genotype data obtained from a cohort of approximately 8.9 million donors (NMDP/Be The Match registry; supplementary Table 1). We analysed a total of 25,128 unique haplotypes to enable 99% coverage of each population which in turn required the analysis of 803 HLA class II molecules (see supplementary information). Supplementary Figure 1 depicts the number of distinct HLA-DRB1, -DRB345 alleles and HLA-DQA1/DQB1, DPA1/DPB1 heterodimers examined and associated haplotype coverage in each population. To confirm the robustness of our observations at the individual level, we analysed multiple replicate genotype samples and demonstrate with replicate sets of simulated genotypes of 10,000 individuals that HLA diversity, as measured by the alpha parameter fits to the power law distribution, was stable, and the number of unique haplotypes needed to reach 95% cumulative frequency was also stable across replicates (supplementary Table 6).

Immunogenic T-cell epitopes were identified using the full-length reference SARS-CoV-2 sequence^20^ with 9,744 amino acids, which was subdivided into the four structural proteins (E, M, N, and S) and 7 additional open reading frames encoding non-structural proteins (NSP1-16) and accessory proteins (proteins 3a, 6, 7a, 7b, 8 and 10). A sliding window of 15 amino acids length was used and, to minimise redundancy, peptides were only counted towards totals if the HLA class II binding core was unique. To evaluate peptide binding and stable display by HLA class II molecules we employed NetMHCIIpan-4.0 which outputs predicted peptide-HLA binding affinity (IC50) in nanomolar units and percentile rank of binding affinity compared to a set of 100,000 random natural peptides. The percentile rank enables incorporation of information from the biological antigen presentation pathway in addition to the peptide-MHC binding event^18, 21^. We aimed to detect strongly binding peptides and, therefore, elected to use a threshold of ≤2% percentile rank (default parameter for HLA class II strong peptide binding) combined with an affinity threshold ≤500nM^18^. To further increase the stringency of criteria for prediction of peptide binding, such that we maximise precision and reduce the rate of false positive peptides (accepting we may not recall all possible peptides that could be displayed), we also explored a model involving a threshold of ≤0.5% percentage rank and ≤500nM binding affinity. Finally, we examined a binding affinity threshold of ≤50nM, as previously suggested^15^. To validate the computational models we analysed publicly available datasets of experimentally determined, immunogenic SARS-CoV-2 peptides. We focused on the two largest datasets recently described by Snyder et al^22, 23^ (336 HLA class II peptides) and by Mateus et al^24^ (135 HLA class II peptides) that contain peptides from the entire SARS-CoV-2 proteome. We also examined two relatively small datasets encompassing 9 nucleocapsid and 25 structural protein-derived (Spike, nucleocapsid or membrane) peptides^25, 26^. In the aforementioned datasets, the HLA restriction was not known and validation focused on positive identification of experimental peptides by the computational models (supplementary information). Finally, we used an HLA-DRB1*04:01 restricted peptide dataset determined using an *in vitro* peptide-HLA stability assay^27^. Our analyses showed that scoring peptide binding based on a combination of ≤2% percentile rank and ≤500nM binding affinity achieved the best true positive rate (sensitivity) for predicting experimentally derived SARS-CoV-2 peptides (supplementary Table 2). Similarly, in the HLA restricted dataset by Prachar et al, the combined ≤2% percentile rank and ≤500nM binding affinity threshold had an AUROC of 0.85 and provided the best combination of precision and specificity in classifying stable peptide binders compared to alternative scoring methods (supplementary Table 3). It was also notable that using a 50nM threshold, without accounting for binding characteristics of natural peptides, resulted in 46% of HLA alleles examined lacking presentation of any SARS-CoV-2 peptides and a bias towards HLA-DR as the major SARS-CoV-2 peptide presenting molecules (supplementary Figure 2). Nevertheless, we have confirmed key aspects of our analyses using both a ≤0.5% percentage rank in combination with a ≤500nM peptide binding affinity threshold, and using a ≤50nM peptide binding affinity threshold alone. Overall, a total of 9,590 15-mer peptides, derived from all 11 SARS-CoV-2 genes, were examined and 4,289 peptides were predicted to bind strongly according to our defined criteria to at least one HLA class II molecule.

### SARS-CoV-2 viral proteome presentation at the molecular HLA class II level

We first examined presentation of viral epitopes by all HLA class II molecules contained in haplotypes representing 99% of the four major broad ethnic populations. Assessment of presentation capacity at the entire viral proteome level (Figure 1A) showed that the majority of HLA class II molecules are capable of presenting SARS-CoV-2 peptides, albeit with significant variability. HLA-DR alleles have the highest viral peptide presentation capacity followed by HLA-DP and, to a significantly lower extent, HLA-DQ molecules. Notably, certain common individual HLA molecules were predicted to have very limited ability to present viral peptides, including DQA1*03:01~DQB1*02:01 (no peptide presentation from any protein within the SARS-CoV-2 proteome; with frequency of 3.2% within AFA haplotypes, <0.01% within API haplotypes, 0.1% within CAU haplotypes and 0.5% within HIS haplotypes), DRB1*03:02 (two peptides presented in total from the entire proteome; with frequency of 6.3% within AFA haplotypes, 0.01% within API haplotypes, 0.04% within CAU haplotypes and 1.0% within HIS haplotypes). Consistent with its known high immunogenicity^28, 29, 30, 31^, Spike protein derived peptides showed strong binding for the majority of HLA class II molecules although presentation capacity again varied and was lowest (no peptides presented) for relatively common alleles such as DQA1*01:01~DQB1*05:03, with a frequency of 1.9-5.4% in each of the four broad population groups (Figure 1B). This observation was more prominent for Nucleocapsid derived peptides where strong binding was predicted to be absent in 117 out of 306 HLA molecules found in 99% of haplotypes in the four broad ethnic populations (mostly reflecting HLA-DP molecules and to some extent -DQ molecules; Figure 1C). Similar findings were noted for the relatively small Membrane and Envelope structural proteins, with the latter predicted to be non-immunogenic for the majority of common HLA class II molecules (Figures 1D-E). For non-structural proteins, HLA class II presentation capacity was variable and dependent upon protein amino acid sequence length (data not shown).

**Figure 1.**
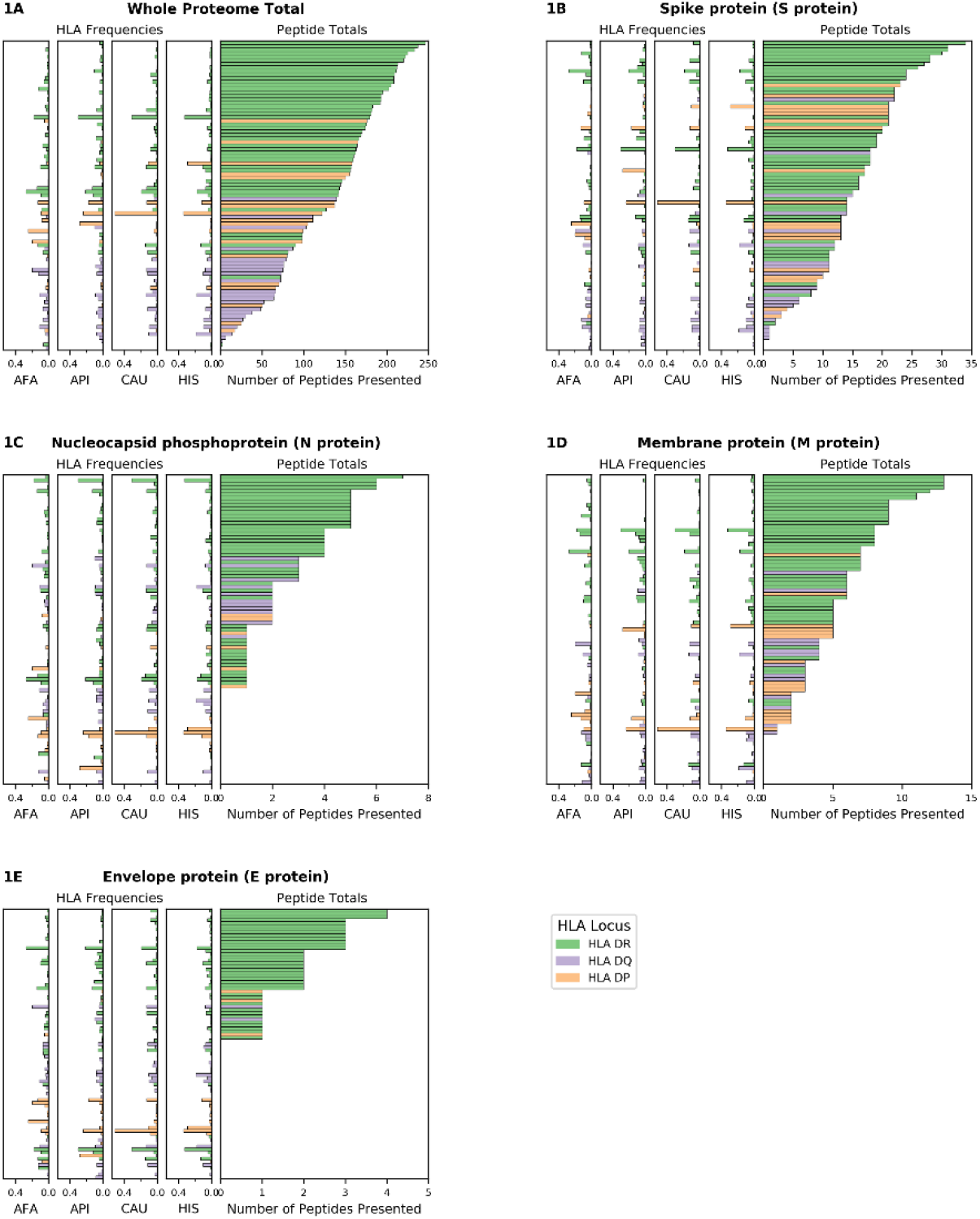
SARS-CoV-2 viral proteome presentation at the molecular HLA class II level. The number of viral peptides presented by individual HLA is shown for A. the entire SARS-CoV-2 proteome; B. Spike protein (S protein); C. Nucleocapsid protein (N protein); D. Membrane protein (M protein); and E. Envelope protein (E protein). The HLA frequency in four broad ethnic populations is shown (AFA: African Americans, API: Asian and Pacific Islanders, CAU: Caucasians, HIS: Hispanics). Bars are coloured according to HLA locus, as shown. HLA were included in the figure if they had a frequency of ≥ 1% in any haplotype distribution from the four broad population groups.

### SARS-CoV-2 proteome immunogenicity at the population level

The capacity of individuals to present viral peptides for recognition by CD4^+^ T-cells depends on the composition of their HLA class II alleles in their inherited haplotypes. Given variation in population-specific HLA haplotype frequencies, we hypothesised that potential differences in SARS-CoV-2 proteome immunogenicity (as reflected by HLA class II peptide presentation) at the population level may reflect disparities in capacity for effective anti-SARS-CoV-2 immunity which could in turn influence response to vaccination or underpin observed variability in COVID-19 clinical outcomes among different ethnic populations. To investigate this hypothesis, we examined viral proteome presentation by HLA class II, accounting for the distribution of HLA haplotypes in a population. This analysis showed that the overall capacity for HLA peptide presentation, at the whole SARS-CoV-2 proteome level, among the four broad ethnic populations examined was remarkably similar (Figure 2A). This was also the case considering T-cell epitopes from each SARS-CoV-2 protein (supplementary Table 4), suggesting that polymorphism at the HLA genomic region is unlikely to underpin potential differences in immune responses and, thus, in clinical outcomes among ethnic groups. It was notable, however, that within ethnic populations the capacity of individual HLA haplotypes to present viral peptides varied widely, with the top 5% of haplotypes predicted to present between 497 and 591 peptides from the entire viral proteome (such as DRB1*04:01~DRB4*01:01~DQA1*03:01~DQB1*03:01~DPA1*01:03~DPB1*04:01 which represents 1% of AFA haplotypes, 0.14% of API haplotypes, 3.9% of CAU haplotypes and 0.9% of HIS haplotypes, predicted to present 506 SARS-CoV-2 peptides) as opposed to 5% of haplotypes at the opposite end of the spectrum predicted to present approximately 316 viral peptides or fewer (e.g. DRB1*03:01~DRB3*01:01~DQA1*05:01~DQB1*02:01~DPA1*02:01~DPB1*01:01 which represents 0.25% of AFA haplotypes, 0.02% of API haplotypes, 3.5% of CAU haplotypes and 1.4% of HIS haplotypes, predicted to present 258 peptides). This observation suggests that individual capacity to mount CD4^+^ T-cell immune responses against SARS-CoV-2 is not uniform and is likely dependent on HLA phenotype. Similar inter-individual variability was noted for peptide presentation derived from structural and from non-structural proteins as well as for distinct SARS-CoV-2 proteins examined (Figure 2B-E and Supplementary Figure 3). Recent experimental studies suggested that up to 70% of CD4^+^ T-cell responses against SARS-CoV-2 target the Spike, Membrane and Nucleocapsid antigens^11^; our analysis showed the predicted CD4^+^ T-cell response to these structural proteins is highly variable at the haplotype level (Figure 2B-D). In agreement with the observed immunogenicity of Spike protein in experimental studies^28, 29, 30, 31^, relatively high numbers of Spike-derived peptides were predicted to be recognised both within and across ethnic populations (Figure 2B). In contrast, our analysis suggests that on average 13.9% of HLA haplotypes within each population have low capacity (≤2 peptides) to present Nucleocapsid-derived peptides (e.g. DRB1*03:02~DRB3*01:01~DQA1*04:01~DQB1*04:02~DPA1*02:02~DPB1*01:01 which presents one peptide, and accounts for 4.98% of haplotypes found in AFA populations and 0.44% in HIS populations, although it is rare in API and CAU populations; Figure 2C). This inter-individual variability may, in part, account for the heterogeneity in the presence and magnitude of CD4^+^ T-cell and antibody responses against the Nucleocapsid protein noted in recent COVID-19 studies^11, 32, 33^. Among non-structural viral proteins, our analysis suggested NSP3, NSP4 and NSP12 as the most immunogenic, in part reflecting their size, in each population (supplementary Figure 3). The above noted similarity in overall capacity for SARS-CoV-2 peptide presentation among different populations was also observed at different (more stringent) thresholds for HLA-peptide binding, albeit with even higher individual variability within ethnic populations (500nM binding affinity and ≤0.5% percentage rank or ≤50nM binding affinity; supplementary Figure 4).

**Figure 2.**
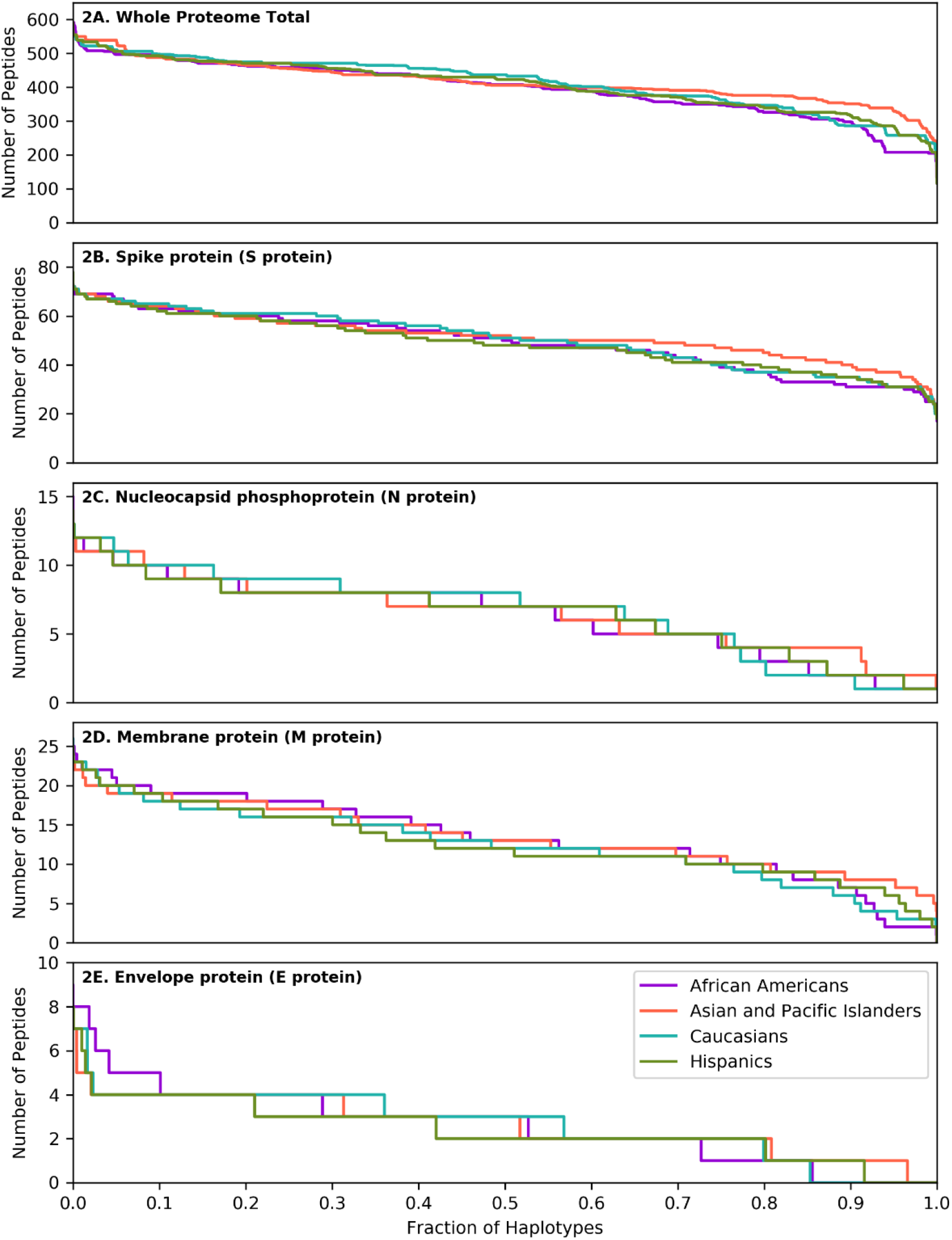
SARS-CoV-2 derived peptide presentation at the HLA haplotype level. Panels depict SARS-CoV-2 peptide presentation by HLA class II haplotypes representing 99% of total haplotypes within four broad population groups (African Americans, Asian Pacific Islanders, Caucasians and Hispanics). A. Whole Proteome. B. Spike protein (S protein). C. Nucleocapsid protein (N protein). D. Membrane protein (M protein). E. Envelope protein (E protein). The width of each step in the curves is proportional to the relative frequency of a specific HLA haplotype.

To further explore the consequences of the above observations at the individual level, and given that every individual expresses two HLA haplotypes, we generated 10 replicates each of random genotype datasets encompassing 1,000, 5,000 and 10,000 simulated individuals for each population, as described in the methods. Populations encompassing 10,000 individuals achieved >95% cumulative HLA haplotype coverage in every population. As shown in Figure 3, this analysis confirmed equivalent HLA class II presentation of SARS-CoV-2 peptides across all four broad ethnic populations, both at the entire viral proteome level and for individual proteins (supplementary Table 4). As noted for the HLA haplotype analysis, there was wide inter-individual variability in predicted potential for CD4^+^ T-cell immune responses, according to HLA genotype. We noted significant, but variable, capacity for T-cell reactivity against the entire Spike glycoprotein across individuals, whereas reactivity against the Nucleocapsid protein was predicted to be weaker for 10% of individuals in each population, on average (range 7.8-13.3%, as defined by HLA presentation of less than 5 nucleocapsid peptides). These observations were confirmed (but, again, inter-individual variability was higher) using more stringent thresholds for defining HLA class II peptide presentation (data not shown). In further analysis, we considered immune reactivity against the Receptor Binding Domain (RBD) of Spike glycoprotein, as it represents a proposed target of coronavirus subunit vaccines currently in clinical trials^34, 35, 36^. Although, overall, there was no significant difference in predicted CD4^+^ T-cell reactivity at the population level, there were notable differences at the individual level (both based on HLA haplotype and on genotype analyses) with wide variation in predicted RBD specific peptide presentation (Figure 3F). The above analyses were consistent irrespective of the size of the population sampled and among the 10 replicates at each population size (Kruskal-Wallis test p-value >0.05 for peptide comparisons at the population level).

**Figure 3.**
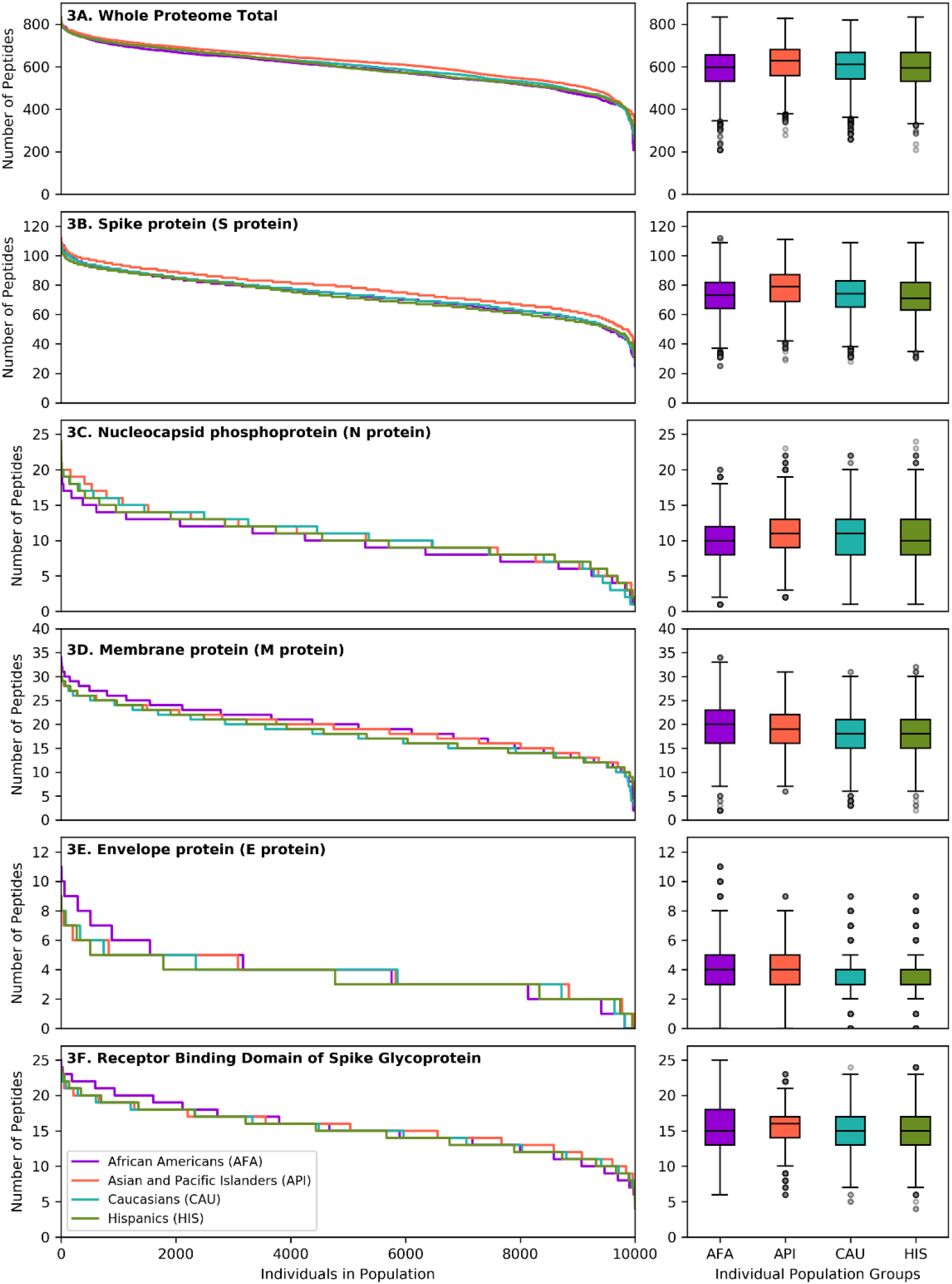
SARS-CoV-2 derived peptide presentation at the HLA genotype level (broad population groups). Panels depict the number of SARS-CoV-2 peptides presented by individual HLA class II genotypes in simulated populations of 10,000 individuals for four broad population groups (African Americans, Asian Pacific Islanders, Caucasians and Hispanics). A. Whole Proteome. B. Spike protein (S protein). C. Nucleocapsid protein (N protein). D. Membrane protein (M protein). E. Envelope protein (E protein). F. Receptor Binding Domain of Spike protein. The width of each step in the curves is proportional to the relative frequency of a specific HLA genotype. The boxplot charts depict the median, interquartile range (box) and range (whiskers - excluding outliers) for the number of viral peptides presented at the population level for each of the above ethnic groups.

We next examined HLA class II presentation of SARS-CoV-2 peptides for a further 21 detailed population subgroups, as described above (Figure 4/supplementary Table 4). Overall, analysis of the entire viral proteome identified similar capacity for HLA class II viral peptide presentation across population subgroups. This was also the case for structural proteins, including Spike and RBD, mirroring the findings above for the broad population groups. Although we did not identify SARS-CoV-2 vulnerability of particular populations at the HLA level, again we observed inter-individual variation in predicted cellular immunity within ethnic groups. This was reflected in the range of predicted viral peptide presentation within simulated populations of 10,000 individuals including 24-112 for Spike, 4-25 for RBD, 1-24 for Nucleocapsid and 208-854 for the entire proteome (Supplementary Table 4).

**Figure 4.**
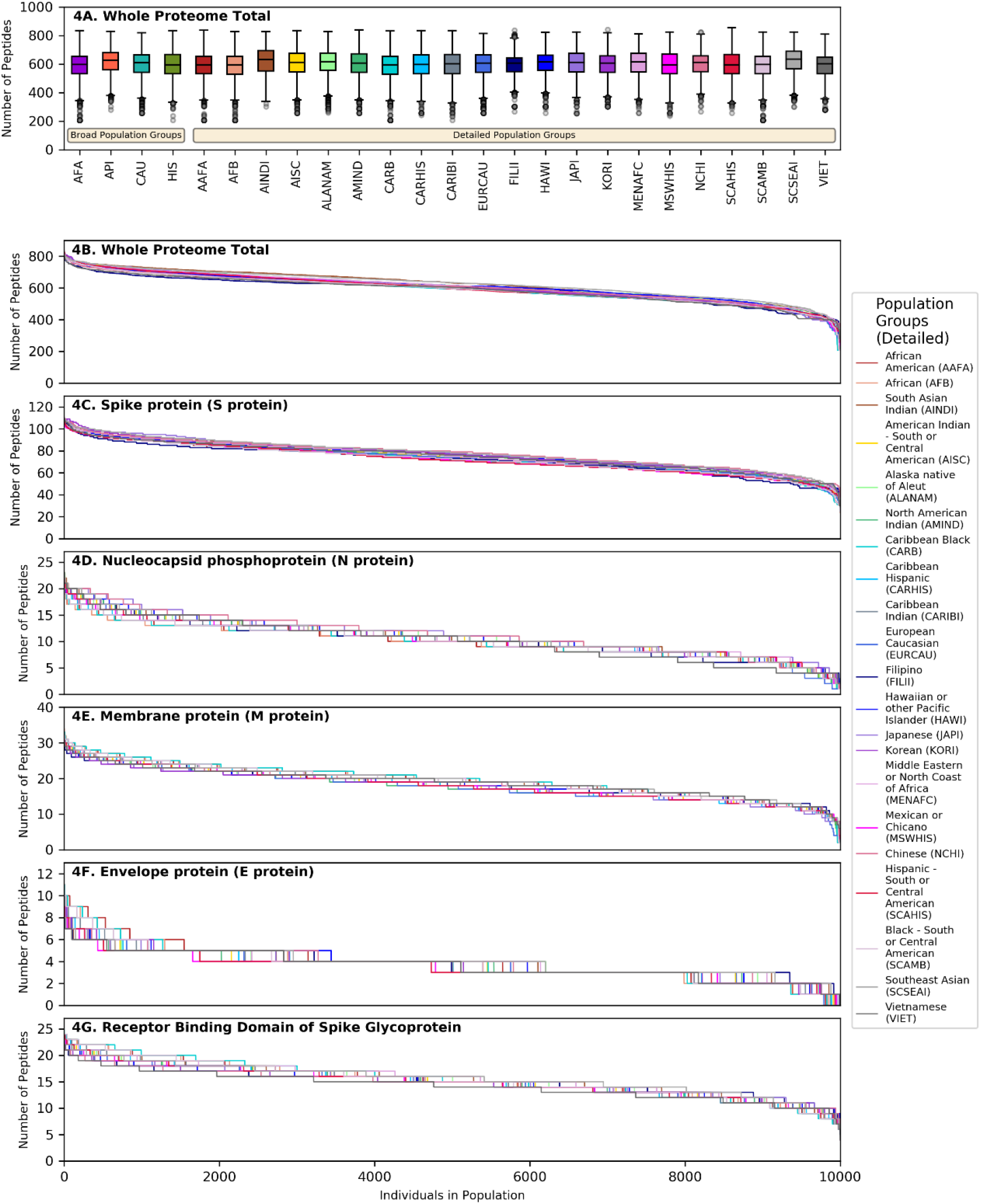
SARS-CoV-2 derived peptide presentation at the HLA genotype level (detailed population subgroups). The top panel (A) shows boxplot charts depicting the median, interquartile range (box) and range (whiskers - excluding outliers) for the number of viral peptides (from the entire SARS-CoV-2 proteome) presented at the population level for each of 4 broad population groups and 21 detailed ethnic population subgroups (shown on the right inset). Panels B-G depict the number of SARS-CoV-2 peptides presented by individual HLA class II genotypes in simulated populations of 10,000 individuals for each of the 21 population subgroups. B. Whole Proteome. C. Spike protein (S protein). D. Nucleocapsid phosphoprotein (N protein). E. Membrane protein (M protein). F. Envelope protein (E protein). G. Receptor Binding Domain of Spike protein.

### Immunogenicity maps of SARS-CoV-2 proteome at the population level

The effectiveness of peptide and subunit vaccine formulations against SARS-CoV-2 depends on robust presentation by individual HLA class II molecules and, therefore, investigated vaccines should account for linkage disequilibrium and HLA haplotype frequencies in different ethnic populations to achieve universal coverage. Capitalising on extensive HLA haplotype frequency data from the NMDP / Be The Match registry, we generated maps of SARS-CoV-2 immunogenicity for the entire viral proteome. This analysis showed that each viral protein contains immunogenic peptide segments with variable degree of population coverage which is, on the whole, similar across different populations for a given protein region, as well as peptide segments of variable length that are non-immunogenic in any population (Figure 5). Similar observations were made for coronavirus subunit components that are being investigated as potential vaccines with several immunogenic peptides predicted to achieve universal coverage across population groups (Figure 5). This was particularly evident for the Spike protein underlying its inherent immunogenicity and its potential as vaccination target. Supplementary Table 5, depicts SARS-CoV-2 immunogenic peptide segments predicted to cover over 90% of HLA genotypic variation in every broad ethnic group examined.

**Figure 5.**
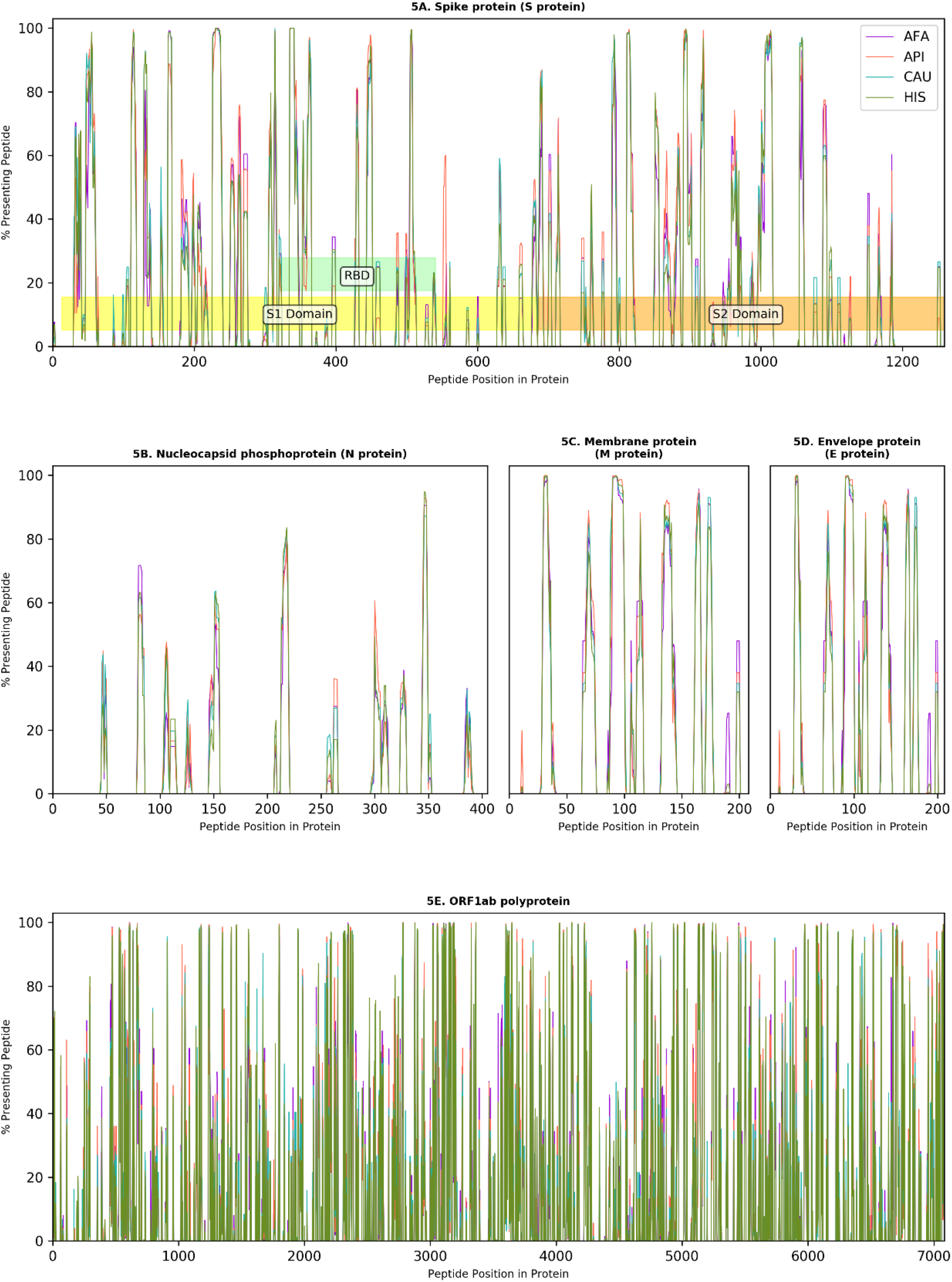
Immunogenicity maps of SARS-CoV-2 proteome at the population level. The panels depict the percentage of HLA class II genotypes, within populations of 10,000 individuals (AFA: African Americans, API: Asian Pacific Islanders, CAU: Caucasians and HIS: Hispanics), that are predicted to present individual SARS-CoV-2 peptides from A. Spike protein (S protein), S1, S2 and Receptor Binding Domain (RBD) are also shown; B. Nucleocapsid protein (N protein); C. Membrane protein (M protein); D. Envelope protein (E protein); and E. Open Reading Frame (ORF) 1ab polyprotein.

## Discussion

Recognition of SARS-CoV-2 peptides in the context of HLA class II molecules is essential for CD4^+^ T-cell activation and proliferation which, in turn, orchestrate the development of effector cellular (CD8^+^ T-cell) and humoral adaptive immune responses after viral infection and after vaccination. In this computational study, we investigated the role of HLA on SARS-CoV-2 immunogenicity at the individual and at the population level, considering population-specific HLA allele, haplotype, and genotype frequencies. We accounted for genetic polymorphism and for HLA linkage disequilibrium in twenty five ethnic populations by capitalising on, to our knowledge, the most extensive HLA haplotype frequency information to date to predict SARS-CoV-2 specific CD4^+^ T-cell epitopes covering the entire viral proteome. We find that the overall capacity for anti SARS-CoV-2 cellular immunity according to HLA class II genotype is similar at the population level across all ethnic groups examined. However, we identify wide inter-individual variability in predicted CD4^+^ T-cell reactivity against every SARS-CoV-2 protein according to expressed HLA genotype. We predict robust immune reactivity against the SARS-CoV-2 Spike protein, the basis for the majority of current vaccination efforts, both at the population and at the individual level regardless of population origin. Finally, we provide comprehensive maps of SARS-CoV-2 proteome immunogenicity accounting for population coverage in ethnic groups.

Several recent studies have examined the cellular immune response to SARS-CoV-2 and revealed strong associations between the T-cell response and COVID-19 severity^29, 37^. Although this relationship is complex to untangle when the peripheral T-cell repertoire is sampled during the acute phase, it is notable that SARS-CoV-2 specific CD4^+^ T-cells have been associated with lessened COVID-19 severity and that high frequency of Spike-specific CD4^+^ T-cell responses were observed in patients who had recovered from COVID-19^11, 29, 38, 39^. A coordinated and regulated response involving all branches of adaptive immunity (CD4^+^, CD8^+^ and antibody responses) is likely required to reduce COVID-19 severity, with the cellular response being key for both initiating the adaptive response and for controlling the acute infection^29^. Even though neutralising antibody titres are not predictive of disease severity^29, 40^, humoral responses are a key aspect of protective immunity after infection and critical for generating sterilising immunity after vaccination^34, 41^. In this respect, current evidence suggests a strong association between the magnitude of Spike-specific CD4^+^ T-cells and neutralising antibody titres^11, 30, 38^. Finally, the majority of recent literature on anti-SARS-CoV-2 immunity indicates there is a high degree of heterogeneity in the breadth and magnitude of both humoral and cellular responses to SARS-CoV-2 both at the individual patient level and in relation to specific viral proteins^11, 29, 33^. Within this context, the observation that patients with severe COVID-19 had decreased diversity of T-cell responses suggested that recognition of multiple SARS-CoV-2 T-cell epitopes may be required for development of protective immunity after infection or after vaccination^42^.

The above observations on anti-SARS-CoV-2 cellular immunity from experimental studies and the role of HLA in shaping the diversity of the T-cell repertoire^43, 44^, place the findings of our study into context. With regards to susceptibility to COVID-19, we hypothesised that genetic variation at the HLA complex may account for observed differences in clinical outcomes between ethnic groups^2, 45, 46^. We performed a comprehensive analysis of HLA haplotypes and genotypes covering 99% of genetic HLA variation within twenty five ethnic populations and showed that the predicted CD4^+^ T-cell response is overall remarkably similar at the population level both looking at the entire SARS-CoV-2 proteome and for individual viral proteins. This observation is supported by more nuanced recent investigations which show equivalent COVID-19 clinical outcomes after adjustment for potential socioeconomic and clinical confounders^7, 8^. We did, however, find significant inter-individual variability in predicted SARS-CoV-2 proteome immunogenicity according to HLA phenotype. This variability was more pronounced for particular viral proteins, such as nucleocapsid, membrane protein and envelope protein and less evident when the entire viral proteome was considered where, even at the lower end of the spectrum, HLA haplotypes were predicted to present a significant number of CD4^+^ T-cell epitopes. Given the relevance of diversity and magnitude of cellular immunity against SARS-CoV-2, as discussed above, it is tempting to speculate that HLA phenotype might underpin some of the observed inter-individual variability in COVID-19 outcomes, along with more established clinical factors. This might depend on the relative contribution of SARS-CoV-2 proteins to the quality of the immune response. For example, Grifoni et al^11^ have shown that up to 70% of CD4^+^ T-cell responses against SARS-CoV-2 target the Spike, Membrane and Nucleocapsid antigens whereas significant reactivity was also noted against nsp3, nsp4 and ORF8. Despite significant differences between the highest and lowest peptide presenting HLA haplotypes, we noted substantial numbers of Spike-specific CD4^+^ T-cell peptides presented by the majority of HLA haplotypes in all ethnic groups examined; in comparison, inter-individual variability according to HLA haplotype was more pronounced for the remaining of the above, and other, viral proteins. Whether this observation might translate into differential clinical outcomes or levels of protective immunity according to expressed HLA type would need a large clinical study that encompasses a cohort representative of the HLA polymorphism within a particular population. Certainly, evidence supporting an important role of HLA class II in viral immunity has been previously reported^47, 48^. The current SARS-CoV-2 pandemic represents a unique opportunity to address such fundamental questions which have recently started to be explored^49^.

It is also well established that individual response, including efficacy and relative antibody levels, and maintenance of immunity after vaccination varies markedly and this biological variation results from a combination of environmental (such as age, size, sex, comorbid status, ethnicity, and dose and route of vaccine administration) and genetic factors^14, 48^. Non-responsiveness affects approximately 2-10% (and up to 20% following hepatitis B vaccination) of vaccinated healthy individuals^50, 51^. HLA class II haplotype plays a central role in the presentation of vaccine epitopes and is a known genetic risk factor for primary vaccination failure^50, 52, 53^. Given that the majority of current vaccination efforts are focused on generating immunity against the Spike protein of SARS-CoV-2, we calculated the number of Spike-specific CD4^+^ T-cell epitopes according to HLA genotype. Our analysis suggests that immune reactivity against Spike is likely to be robust both at the population level, including all 25 ethnic groups examined, and at the individual level. This finding is now supported experimentally by studies reporting high degree of seroconversion against Spike after natural infection and after vaccination^28, 30, 33, 54, 55, 56, 57, 58, 59^. Nevertheless, we noted inter-individual variation ranging from 112 peptides for the highest presenting HLA class II genotypes to 24 for the lowest. Whether such variation may affect the magnitude and diversity of protective immunity generated after vaccination requires further study but it is notable that variation in the degree of cellular immunity has been reported with a few vaccine formulations^55, 58, 60^.

It is important to acknowledge the limitations of our study. We used a computational approach to predict SARS-CoV-2 peptides presented by HLA class II molecules, however, peptide presentation does not always lead to CD4^+^ T-cell activation; peptide recognition is complex and incompletely understood and is influenced by many factors, including relative expression of individual viral proteins^61^. Nevertheless, NetMHCIIpan-4.0 is an established and validated algorithm for T-cell epitope prediction that has recently been updated resulting in improved performance^21^. Recent computational studies investigating SARS-CoV-2 vaccine immunogenicity have based their approach for T-cell epitope selection exclusively on peptide-HLA binding affinity incorporating different thresholds (e.g. 500nM or 50nM) and identified population coverage gaps in predicted cellular immunity^15, 62, 63, 64^. This approach is affected by inherent bias of certain HLA molecules towards higher or lower mean predicted affinities; thus, we show that the 50nM binding affinity threshold, one of the most commonly used, is heavily biased towards HLA-DR as the main SARS-CoV-2 peptide presenting locus with the majority of HLA-DQ and -DP molecules showing no peptide binding. Accordingly, using a 50nM binding affinity threshold for defining peptide immunogenicity resulted in very wide inter-individual variability in predicted CD4^+^ T-cell reactivity against SARS-CoV-2 proteins (supplementary Figure 4). To overcome this limitation, we incorporated both binding affinity prediction and percentile rank, compared to a set of 100,000 random natural peptides, for epitope selection. Notwithstanding that currently there is limited information on experimentally determined SARS-CoV-2 immunogenic T-cell epitopes and even less information on peptide HLA restriction, we have used available information in the published literature to validate our approach. We focused on prediction of strongly HLA binding peptides and demonstrated that our peptide selection threshold correctly identified the majority of experimentally determined immunogenic viral peptides in the largest published datasets^22, 24^. We also aimed to minimise the rate of false positive peptides and showed, in a limited experimental SARS-CoV-2 peptide dataset with HLA restrictions published by Prachar et al^20^, that our 500nM binding affinity and ≤2% rank threshold for peptide selection, achieves high precision and specificity (1.0 for both). Finally, we confirmed that our observations on SARS-CoV-2 immunogenicity, both in relation to the comparison of population level responses and in relation to inter-individual variability, remained valid when we used more stringent peptide selection criteria (500nM and ≤0.5% rank) and the commonly used 50nM binding affinity threshold.

In conclusion, we present a rigorous immune-informatics approach to evaluate the potential for cellular immunity against SARS-CoV-2 at the population and at the individual level capitalising on, to our knowledge, the most comprehensive assessment of HLA genetic variation to date. Our findings provide important insight on the potential role of HLA polymorphism on development of protective immunity after SARS-CoV-2 infection and after vaccination and a firm basis for further experimental studies in this field.

## Acknowledgments

The research presented in this manuscript has received funding from the NIHR Blood and Transplant Research Unit in Organ Donation and Transplantation at the University of Cambridge, from the NIHR Cambridge Biomedical Research Centre and from an NIHR Fellowship (PDF-2016-09-065, VK). We gratefully acknowledge funding from an MRC Clinical Research Training Fellowship to HCC (MR/S006745/1). ARL acknowledges funding by the Member States of the European Molecular Biology Laboratory (EMBL). We thank the US Registry National Marrow Donor Program / Be The Match and the 38+ million stem cell registry volunteers worldwide. The views expressed are those of the authors and not necessarily those of the National Health Service, the National Institute for Health Research, the Department of Health, or National Health Service Blood and Transplant.

