## Supplementary Information for "Influence of HLA class II polymorphism on predicted cellular immunity against SARS-CoV-2 at the population and individual level"

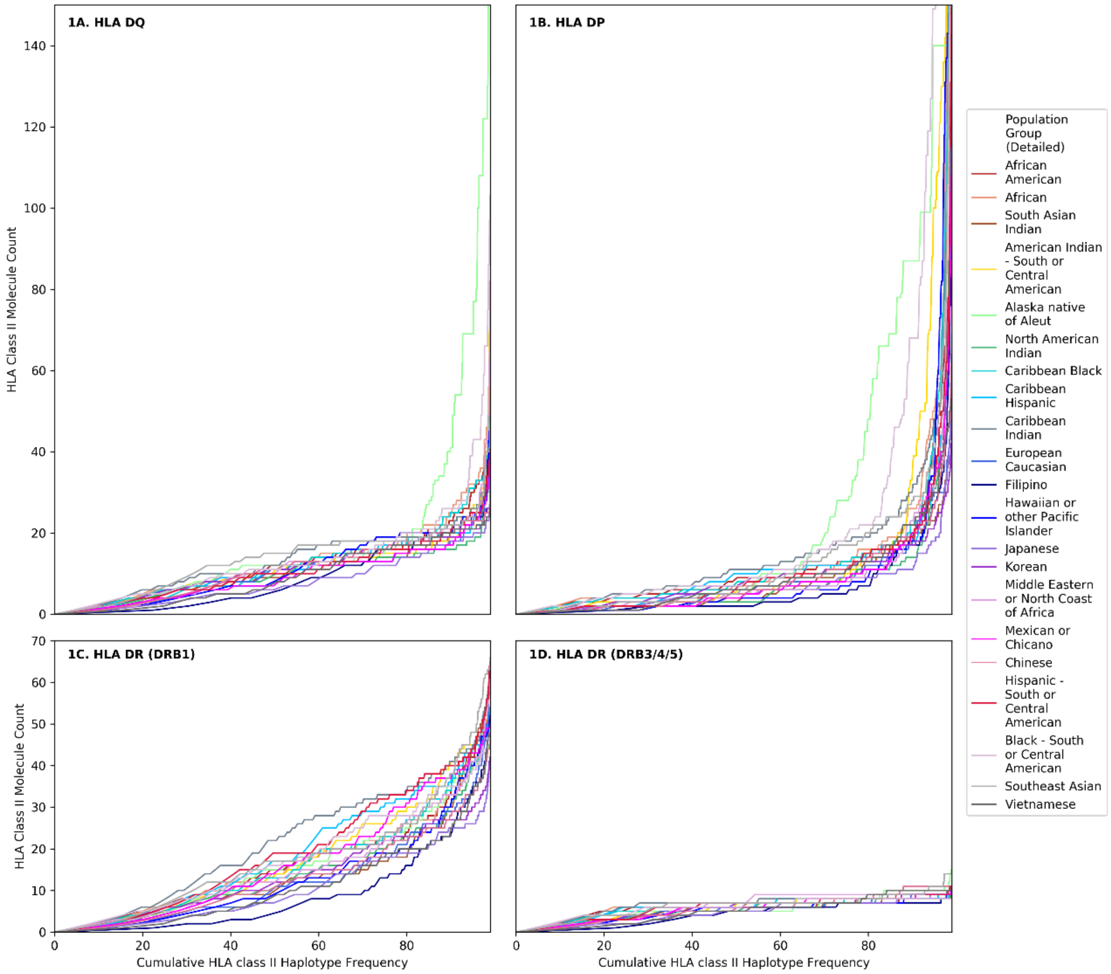


***Supplementary Figure 1. Cumulative HLA class II haplotype frequency.***

The panels show the cumulative HLA class II haplotype frequency (x-axis) and the corresponding HLA Class II molecule count (y-axis) for each HLA class II locus: A. HLA DQ; B. HLA DP; C. HLA DR (DRB1); D. HLA DR (DRB3/4/5). The Figure depicts 21 detailed population subgroups, as shown in the inset. Of the HLA observed, on average 0.52% of haplotypes contained an HLA that was not in the list of 5,620 HLA available to run on NetMHCIIpan-4.0.

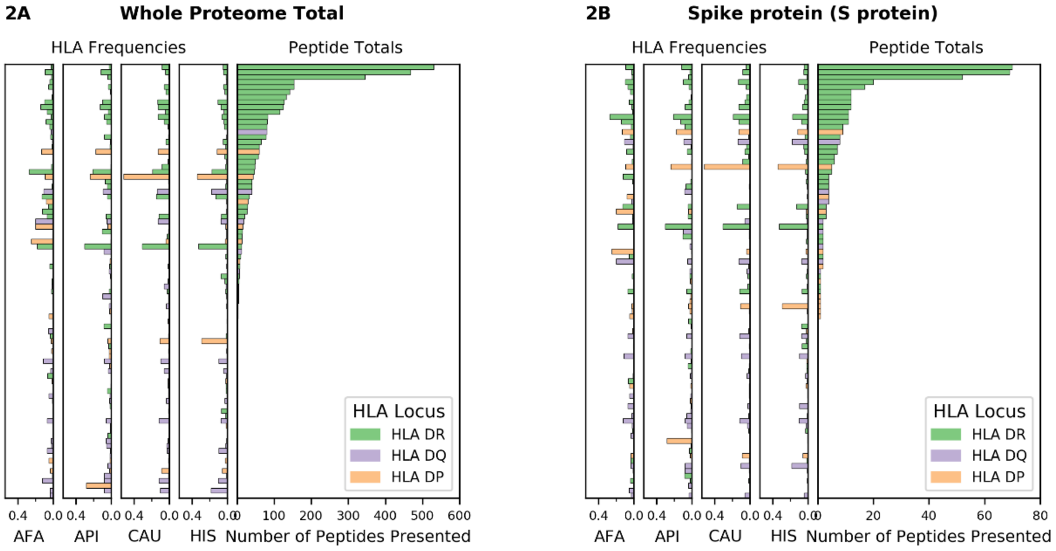


***Supplementary Figure 2. SARS-CoV-2 viral proteome presentation at the molecular HLA class II level (50nM threshold)***

The number of viral peptides presented by individual HLA is shown for A. the entire SARS-CoV-2 proteome; B. Spike protein (S protein). A binding affinity of ≤50nM was used as a threshold to define viral peptide presentation by HLA class II. The HLA frequency in four broad ethnic populations is shown (AFA: African Americans, API: Asian and Pacific Islanders, CAU: Caucasians, HIS: Hispanics). Bars are coloured according to HLA locus, and HLAs were included in the figure if they had a frequency of ≥ 1% in any haplotype distribution from the four broad population groups.

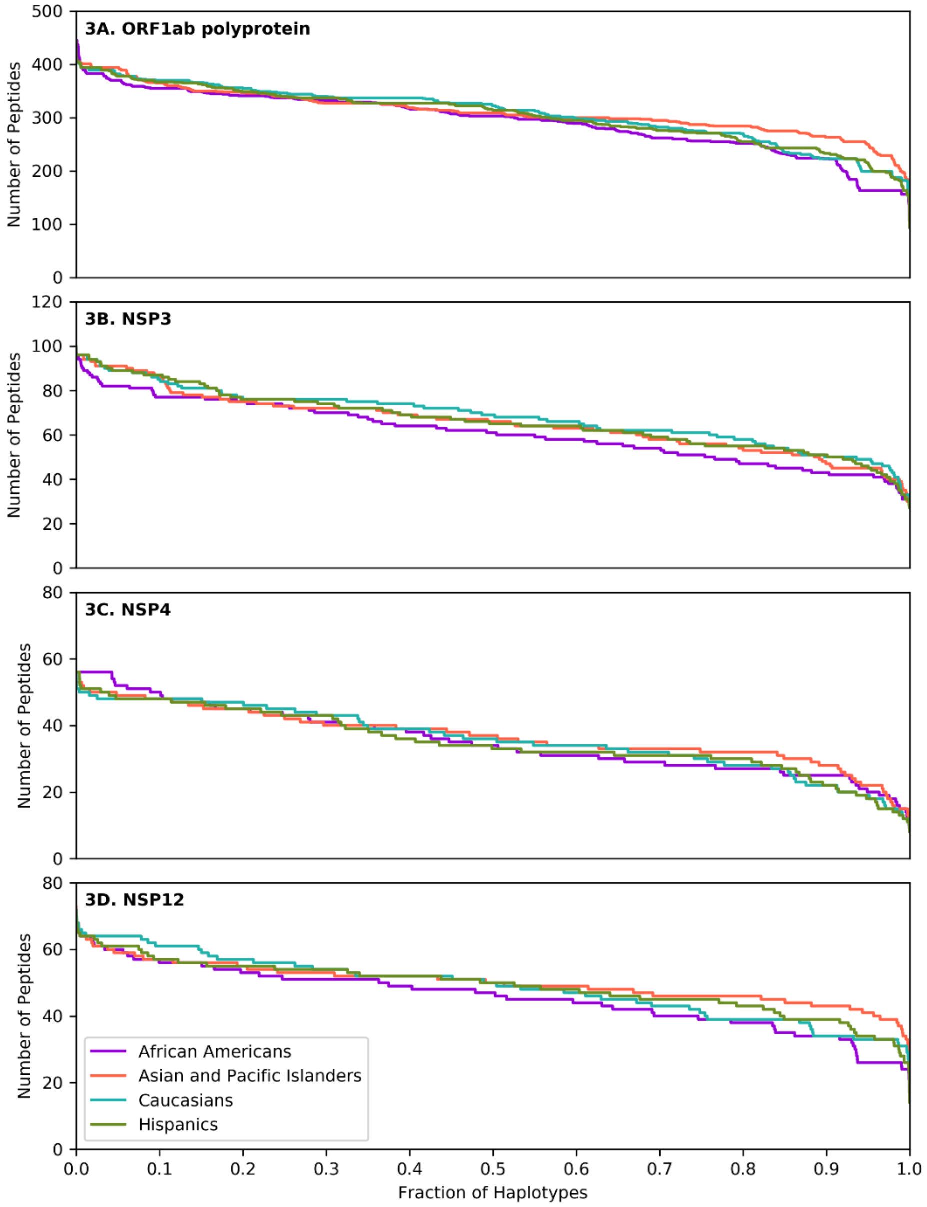


***Supplementary Figure 3. SARS-CoV-2 derived peptide presentation at the HLA haplotype level.***

Panels depict peptide presentation by HLA class II haplotypes representing 99% of total haplotypes within four broad population groups (African Americans, Asian Pacific Islanders, Caucasians and Hispanics). A. ORF1ab polyprotein. B. Non structural protein 3 (NSP3). C. Non structural protein 4 (NSP4). D. Non structural protein 12 (NSP12). The width of each step in the curves is proportional to the relative frequency of a specific HLA haplotype.

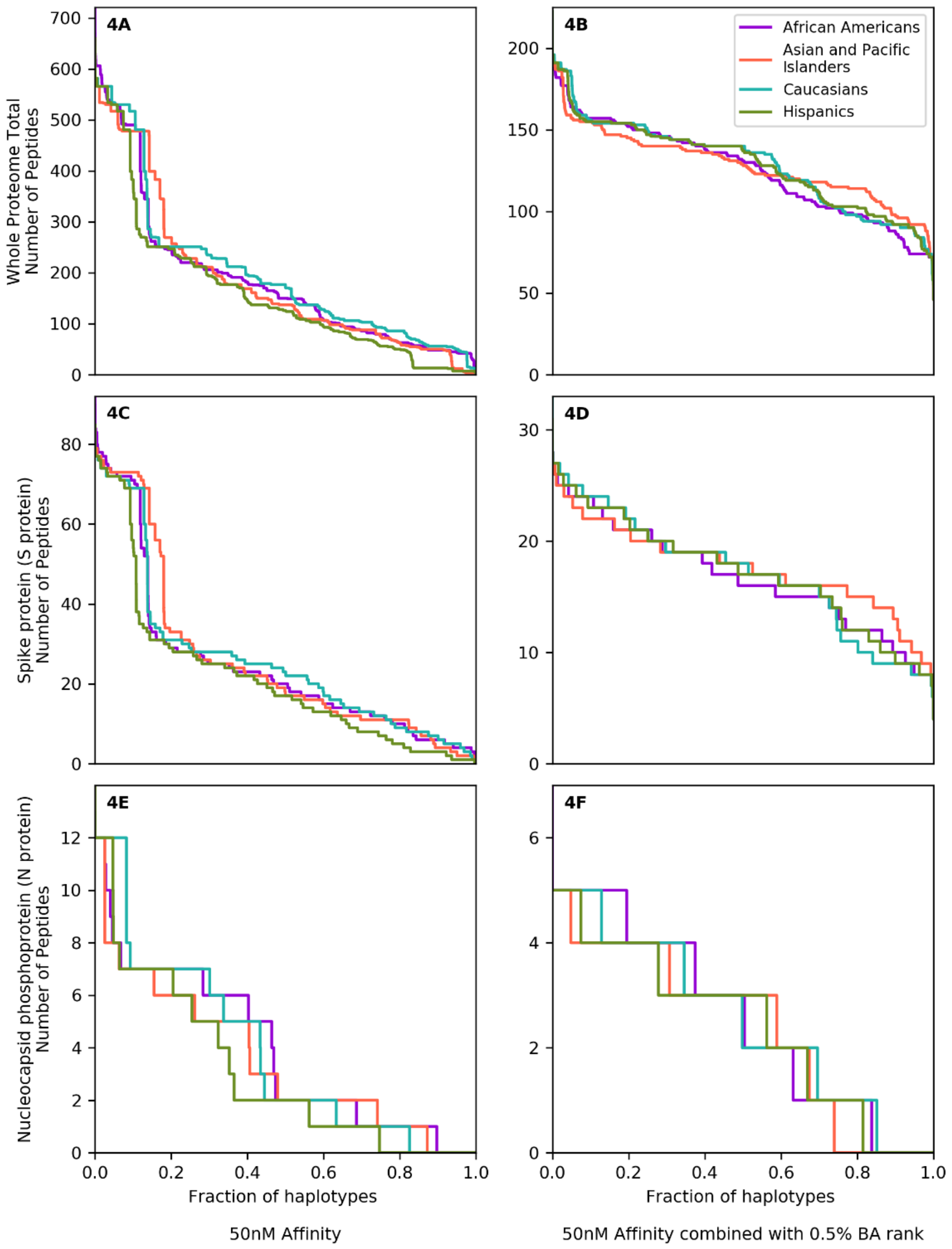


**Supplementary Figure 4. SARS-CoV-2 derived peptide presentation at the HLA haplotype level using two alternative computational models.**

Panels depict SARS-CoV-2 peptide presentation by HLA class II haplotypes representing 99% of total haplotypes within four broad population groups (African Americans, Asian Pacific Islanders, Caucasians and Hispanics). Two computational models utilising different NetMHCIIpan-4.0 HLA-peptide binding thresholds were utilised: in Panels A, C and E (on the left) a ≤50nM HLA-peptide binding affinity threshold was used; in Panels B, D and F (on the right) a combined ≤2% percentile rank and ≤500nM binding affinity thresholds was used. 4A and 4B. Whole Proteome. 4C and 4D. Spike protein (S protein). 4E and 4F. Nucleocapsid phosphoprotein (N protein). The width of each step in the curves is proportional to the relative frequency of a specific HLA haplotype. BA: Binding Affinity.

| | Population Group | Number of Individuals HLA Typed by molecular methods | | | | | | | Number of Individuals HLA Typed at High Resolution | | | | | | | | |
|---|---|---|---|---|---|---|---|---|---|---|---|---|---|---|---|---|---|
| | | HLA-A, HLA-B, HLA-DRB1 | HLA-C | HLA-DRB3/4/5 | HLA-DQA1 | HLA-DQB1 | HLA-DPA1 | HLA-DPB1 | HLA-A | HLA-B | HLA-DRB1 | HLA-C | HLA-DRB345 | HLA-DQA1 | HLA-DQB1 | HLA-DPA1 | HLA-DPB1 |
| Broad Population Groups | African Americans | 871,096 | 430,069 | 362,263 | 22,523 | 317,862 | 20,068 | 218,650 | 317,470 | 388,347 | 483,154 | 262,619 | 106,130 | 21,245 | 269,904 | 17,043 | 214,672 |
| | Asian and Pacific Islanders | 982,748 | 473,379 | 392,138 | 30,424 | 386,893 | 28,213 | 276,378 | 352,285 | 415,956 | 536,974 | 293,601 | 133,884 | 27,388 | 321,615 | 23,093 | 270,626 |
| | Caucasians | 5,613,087 | 3,608,934 | 2,289,796 | 69,216 | 2,911,375 | 64,796 | 2,066,424 | 2,506,083 | 2,622,663 | 3,556,145 | 2,266,283 | 1,108,008 | 63,037 | 2,614,010 | 62,258 | 2,046,765 |
| | Hispanics | 1,357,465 | 755,940 | 551,167 | 18,401 | 603,342 | 15,415 | 441,513 | 547,343 | 626,392 | 808,061 | 472,300 | 249,916 | 17,405 | 534,848 | 14,799 | 437,281 |
| Narrow Population Groups | African American | 654,858 | 323,818 | 263,989 | 17,008 | 229,485 | 15,185 | 160,508 | 234,940 | 293,123 | 359,914 | 194,307 | 68,590 | 16,068 | 194,852 | 12,894 | 157,287 |
| | African | 55,124 | 31,782 | 22,116 | 1,856 | 25,826 | 1,716 | 19,015 | 25,625 | 29,530 | 35,709 | 21,090 | 8,057 | 1,762 | 22,661 | 1,473 | 18,817 |
| | South Asian Indian | 314,032 | 147,304 | 128,349 | 11,783 | 125,093 | 11,219 | 87,494 | 111,602 | 132,018 | 170,142 | 89,879 | 43,361 | 10,291 | 103,936 | 10,615 | 86,937 |
| | American Indian - South or Central American | 10,058 | 5,306 | 3,541 | 279 | 4,117 | 262 | 3,275 | 3,836 | 4,074 | 5,940 | 3,364 | 1,632 | 260 | 3,639 | 253 | 3,243 |
| | Alaska native of Aleut | 4,787 | 3,693 | 3,326 | 77 | 3,445 | 71 | 3,173 | 3,315 | 3,311 | 3,803 | 2,921 | 2,746 | 77 | 3,304 | 68 | 3,162 |
| | North American Indian | 50,818 | 20,548 | 23,748 | 839 | 15,077 | 708 | 9,848 | 13,844 | 15,267 | 21,665 | 12,120 | 5,901 | 753 | 13,019 | 659 | 9,691 |
| | Caribbean black | 58,615 | 34,169 | 21,401 | 1,942 | 25,786 | 1,768 | 17,993 | 24,865 | 29,375 | 37,069 | 20,688 | 6,756 | 1,824 | 22,229 | 1,487 | 17,711 |
| | Caribbean Hispanic | 172,227 | 73,787 | 66,668 | 2,600 | 56,433 | 2,267 | 36,685 | 50,161 | 61,651 | 87,628 | 39,491 | 16,916 | 2,448 | 48,925 | 2,157 | 36,208 |
| | Caribbean Indian | 36,842 | 26,982 | 11,587 | 2,316 | 22,712 | 2,205 | 16,273 | 18,180 | 20,274 | 27,194 | 15,075 | 7,027 | 2,135 | 19,220 | 2,044 | 16,213 |
| | European Caucasian | 2,835,760 | 1,958,391 | 1,149,518 | 41,617 | 1,645,874 | 40,611 | 1,207,951 | 1,426,225 | 1,458,541 | 1,964,329 | 1,277,484 | 647,588 | 38,221 | 1,502,734 | 39,073 | 1,197,733 |
| | Filipino | 86,715 | 49,801 | 33,408 | 2,586 | 35,188 | 2,349 | 26,668 | 35,965 | 42,663 | 48,285 | 29,895 | 13,031 | 2,385 | 29,563 | 1,436 | 25,372 |
| | Hawaiian or other Pacific Islander | 19,871 | 10,048 | 7,598 | 533 | 7,014 | 419 | 4,478 | 6,628 | 7,503 | 9,991 | 5,334 | 2,054 | 502 | 5,541 | 324 | 4,282 |
| | Japanese | 34,855 | 12,221 | 14,363 | 859 | 9,049 | 770 | 5,957 | 8,321 | 9,991 | 14,859 | 7,023 | 2,990 | 706 | 7,108 | 564 | 5,701 |
| | Korean | 120,464 | 50,289 | 52,204 | 2,637 | 40,300 | 2,425 | 29,220 | 38,625 | 43,773 | 58,514 | 32,452 | 16,431 | 2,342 | 33,481 | 1,876 | 28,532 |
| | Middle Eastern or North Coast of Africa | 147,918 | 98,185 | 63,890 | 1,897 | 80,744 | 2,165 | 57,419 | 70,371 | 76,781 | 99,307 | 64,018 | 29,732 | 1,784 | 73,394 | 2,116 | 56,804 |
| | Mexican or Chicano | 456,977 | 238,154 | 208,109 | 4,425 | 187,786 | 3,513 | 140,826 | 174,115 | 200,803 | 245,226 | 154,134 | 87,550 | 4,151 | 168,394 | 3,376 | 139,464 |
| | Chinese | 171,875 | 84,602 | 64,109 | 4,934 | 71,342 | 4,464 | 51,467 | 62,316 | 74,463 | 98,206 | 54,216 | 21,931 | 4,602 | 59,803 | 3,412 | 50,256 |
| | Hispanic - South or Central American | 257,159 | 132,446 | 102,422 | 3,593 | 104,548 | 3,004 | 74,393 | 94,774 | 112,057 | 149,595 | 80,072 | 43,670 | 3,394 | 92,745 | 2,886 | 73,725 |
| | Black - South or Central American | 7,533 | 3,415 | 3,130 | 175 | 2,521 | 162 | 1,780 | 2,527 | 3,105 | 3,899 | 2,088 | 717 | 166 | 2,091 | 141 | 1,738 |
| | Southeast Asian | 53,529 | 31,062 | 19,643 | 1,952 | 26,665 | 1,835 | 20,290 | 23,577 | 26,907 | 34,456 | 20,611 | 9,286 | 1,761 | 23,174 | 1,575 | 20,064 |
| | Vietnamese | 77,865 | 42,572 | 25,914 | 3,431 | 33,269 | 3,254 | 24,833 | 32,020 | 38,905 | 51,663 | 24,480 | 11,058 | 3,279 | 26,842 | 2,057 | 24,025 |

***Supplementary Table 1. National Marrow Donor Program / Be The Match HLA genotype data.*** Number of individuals from different ethnic groups HLA typed by molecular methods and subset of individuals HLA typed at high-resolution. The total number of individuals was 8,931,688, accounting for the fact that many individuals are present in both narrow and broad population groups.

| Peptide Binding Computational Method | MIRA identified T-Cell Epitopes | Cross Reactive SARS-COV-2 T-Cell Epitopes | T-Cell Epitopes post SARS-CoV-2 | T-Cell Epitopes post SARS-CoV-2 |
|---|---|---|---|---|
| | (Snyder et al) | (Mateus et al) | (Le Bert et al) | (Keller et al) |
| ≤ 500nM binding affinity & ≤0.5% BA percentage rank | 225/336 (67%) | 61/135 (45%) | 4/9 (44%) | 5/25 (20%) |
| ≤ 500nM binding affinity & ≤2% BA percentage rank | 324/336 (96%) | 108/135 (80%) | 7/9 (78%) | 15/25 (60%) |
| ≤ 50nM binding affinity | 269/336 (83%) | 83/135 (61%) | 7/9 (78%) | 13/25 (52%) |

***Supplementary Table 2. Classification performance of computational methods for predicting experimentally determined SARS-CoV-2 peptide immunogenicity.*** Evaluation of computational models utilising different NetMHCIIpan-4.0 HLA-peptide binding thresholds on four sets of experimentally determined SARS-CoV-2 $CD4^+$ T-cell epitopes. In the datasets by Snyder et al[1], Mateus et al[2] and Le Bert et al[3], the HLA restriction of immunogenic peptides was not known. For the purpose of this analysis, an experimental peptide was considered to be correctly identified by a given computational method if it was predicted to be presented by ≥10% of HLA haplotypes within a broad population group. For the dataset by Keller et al[4], the HLA type of the responding patient was known. An experimentally determined peptide in this dataset was considered to be correctly identified by a given computational method if it was predicted to be presented by any of the HLA class II of the responding patient. We found that, irrespective of dataset used, the combined ≤2% percentile rank and ≤500nM binding affinity threshold had the highest true positive rate.

| Peptide Binding Computational Method | AUROC | Precision | Sensitivity | Specificity | Accuracy |
|---|---|---|---|---|---|
| ≤ 50nM binding affinity | 0.85 | 0.000 | 0.000 | 1.000 | 0.763 |
| ≤ 500nM binding affinity & ≤2% BA percentage rank | 0.85 | 1.000 | 0.136 | 1.000 | 0.796 |
| ≤ 500nM binding affinity & ≤0.5% BA percentage rank | 0.85 | 0.000 | 0.000 | 1.000 | 0.763 |

***Supplementary Table 3. Classification performance of computational methods for predicting experimentally determined SARS-CoV-2 peptide-HLA binding.*** Evaluation of computational models utilising different NetMHCIIpan-4.0 HLA-peptide binding thresholds using a dataset of HLA-DRB1*04:01 restricted SARS-CoV-2 peptides (n=93) determined using an *in vitro* peptide-HLA stability assay[5]. We found that classifying peptide binders using a combined ≤2% percentile rank and ≤500nM binding affinity threshold maximised precision, sensitivity, specificity and accuracy. BA = Binding Affinity. AUROC = Area Under the Receiver Operator Curve. For AUROC involving models combining two parameters, the predictor variable used was the normalised sum of both parameters.

| | | Spike protein (S protein) | | Nucleocapsid phosphoprotein (N protein) | | Membrane protein (M protein) | | Envelope protein (E protein) | | ORF1ab polyprotein | | Whole Proteome Total | |
|---|---|---|---|---|---|---|---|---|---|---|---|---|---|
| | | Mean (Standard Deviation) | Range | Mean (Standard Deviation) | Range | Mean (Standard Deviation) | Range | Mean (Standard Deviation) | Range | Mean (Standard Deviation) | Range | Mean (Standard Deviation) | Range |
| Broad Population Groups | African Americans | 73.1 (13.2) | 25-112 | 9.9 (3.1) | 1-20 | 19.6 (4.8) | 2-34 | 4.0 (1.8) | 0-11 | 438.7 (66.4) | 163-625 | 591.1 (90.6) | 208-834 |
| | Asian and Pacific Islanders | 77.7 (12.5) | 29-111 | 11.0 (3.6) | 2-23 | 19.1 (4.1) | 6-31 | 3.9 (1.2) | 0-9 | 461.7 (62.2) | 210-620 | 619.2 (84.3) | 279-829 |
| | Caucasians | 73.7 (13.0) | 28-109 | 10.9 (3.6) | 1-22 | 18.0 (4.4) | 3-31 | 3.8 (1.3) | 0-9 | 449.2 (64.6) | 188-623 | 602.4 (88.0) | 257-820 |
| | Hispanics | 72.1 (13.0) | 30-109 | 10.6 (3.2) | 1-24 | 18.3 (4.4) | 2-32 | 3.5 (1.2) | 0-9 | 446.3 (66.4) | 163-629 | 596.6 (89.0) | 208-834 |
| Detailed Population Groups | African American | 73.0 (13.1) | 25-112 | 9.9 (3.1) | 1-20 | 19.6 (4.7) | 2-34 | 3.9 (1.8) | 0-11 | 437.9 (65.6) | 163-636 | 590.1 (89.4) | 208-837 |
| | African | 73.3 (13.2) | 24-112 | 9.9 (3.1) | 1-22 | 19.7 (4.7) | 2-35 | 3.8 (1.7) | 0-11 | 437.4 (65.9) | 163-628 | 589.3 (89.6) | 208-828 |
| | South Asian Indian | 76.2 (13.4) | 29-107 | 10.9 (3.6) | 2-23 | 19.3 (4.3) | 6-30 | 3.9 (1.1) | 0-8 | 463.1 (66.0) | 233-612 | 621.8 (89.7) | 303-827 |
| | American Indian - South or Central American | 74.0 (13.0) | 27-112 | 10.8 (3.6) | 1-24 | 18.3 (4.5) | 3-33 | 3.7 (1.3) | 0-10 | 453.6 (65.0) | 199-627 | 607.3 (88.1) | 258-835 |
| | Alaska native of Aleut | 75.2 (12.6) | 27-111 | 10.9 (3.6) | 1-24 | 18.3 (4.2) | 3-32 | 3.8 (1.3) | 0-9 | 456.0 (62.8) | 204-616 | 610.8 (85.2) | 259-827 |
| | North American Indian | 73.5 (12.9) | 27-111 | 10.8 (3.6) | 1-23 | 17.9 (4.3) | 3-31 | 3.7 (1.2) | 0-9 | 451.5 (64.9) | 199-630 | 604.5 (87.7) | 258-839 |
| | Caribbean black | 73.0 (13.4) | 25-112 | 9.9 (3.1) | 1-22 | 19.7 (4.8) | 2-34 | 3.8 (1.8) | 0-12 | 437.0 (66.8) | 163-624 | 589.1 (91.3) | 208-832 |
| | Caribbean Hispanic | 72.5 (13.0) | 25-112 | 10.5 (3.2) | 1-23 | 18.4 (4.7) | 3-32 | 3.6 (1.4) | 0-11 | 444.2 (67.2) | 184-624 | 595.1 (90.3) | 241-834 |
| | Caribbean Indian | 72.8 (13.1) | 27-112 | 10.6 (3.2) | 1-23 | 18.7 (4.7) | 2-33 | 3.7 (1.5) | 0-11 | 445.5 (68.3) | 163-629 | 597.0 (91.9) | 208-834 |
| | European Caucasian | 73.5 (13.1) | 27-108 | 10.8 (3.7) | 1-23 | 17.8 (4.5) | 3-31 | 3.7 (1.3) | 0-10 | 447.7 (65.4) | 199-610 | 600.2 (89.4) | 258-816 |
| | Filipino | 71.9 (12.8) | 38-109 | 10.3 (3.4) | 2-21 | 18.6 (3.9) | 8-32 | 3.8 (1.1) | 0-8 | 439.6 (58.4) | 197-620 | 589.0 (79.2) | 267-837 |
| | Hawaiian or other Pacific Islander | 76.0 (12.0) | 35-111 | 11.1 (3.6) | 1-21 | 18.6 (4.0) | 3-33 | 3.9 (1.4) | 0-9 | 453.4 (61.9) | 199-612 | 606.9 (83.8) | 258-823 |
| | Japanese | 76.9 (13.7) | 34-110 | 11.4 (3.7) | 2-22 | 18.0 (4.3) | 8-31 | 3.7 (1.4) | 0-9 | 450.3 (67.8) | 188-613 | 603.6 (93.3) | 254-826 |
| | Korean | 76.6 (13.0) | 32-110 | 10.7 (3.5) | 2-22 | 18.0 (4.0) | 7-31 | 3.7 (1.5) | 0-9 | 446.4 (63.2) | 222-632 | 597.5 (85.7) | 302-842 |
| | Middle Eastern or North Coast of Africa | 75.1 (12.9) | 31-109 | 10.7 (3.3) | 1-22 | 19.3 (4.3) | 3-32 | 3.8 (1.2) | 0-9 | 453.7 (63.3) | 199-604 | 609.1 (85.6) | 258-812 |
| | Mexican or Chicano | 72.1 (13.3) | 27-111 | 10.5 (3.3) | 1-24 | 18.3 (4.4) | 3-32 | 3.5 (1.2) | 0-8 | 446.9 (68.0) | 182-624 | 597.0 (91.2) | 236-826 |
| | Chinese | 77.6 (12.3) | 37-111 | 11.3 (4.0) | 2-21 | 18.9 (4.0) | 7-31 | 3.9 (1.4) | 0-9 | 447.3 (60.6) | 197-610 | 599.7 (82.2) | 267-824 |
| | Hispanic - South or Central American | 72.0 (13.2) | 31-108 | 10.6 (3.3) | 1-24 | 18.2 (4.4) | 3-33 | 3.5 (1.2) | 0-9 | 446.3 (67.2) | 199-635 | 596.7 (90.2) | 258-854 |
| | Black - South or Central American | 72.4 (13.3) | 25-111 | 10.2 (3.1) | 1-22 | 19.2 (4.7) | 2-34 | 3.8 (1.6) | 0-11 | 439.5 (64.5) | 163-624 | 591.0 (88.0) | 208-826 |
| | Southeast Asian | 77.6 (12.9) | 29-112 | 11.0 (3.6) | 2-24 | 19.5 (4.1) | 6-32 | 3.9 (1.1) | 0-8 | 465.4 (62.1) | 225-610 | 624.5 (84.3) | 300-826 |
| | Vietnamese | 74.9 (12.4) | 30-112 | 10.2 (4.0) | 2-22 | 19.0 (3.8) | 6-31 | 3.7 (1.2) | 0-9 | 439.0 (62.5) | 210-600 | 586.4 (84.4) | 279-810 |

***Supplementary Table 4. SARS-CoV-2 derived peptide presentation at the population level.*** Descriptive statistics (number of predicted peptides presented, standard deviation and range) based on analyses of random HLA genotype datasets encompassing 10,000 simulated individuals from four broad population groups and twenty one detailed population subgroups. This is shown for Spike protein (S protein), Nucleocapsid phosphoprotein (N protein), Membrane protein (M protein), Envelope protein (E protein), ORF1ab polyprotein and the entire SARS-CoV-19 proteome.

***Supplementary Table 5. Highly immunogenic SARS-CoV-2 peptide segments.*** The Table depicts immunogenic SARS-CoV-2 peptide segments predicted to cover over 90% of HLA genotypic variation (based on analyses of 10,000 simulated individuals for each population) in all of four major broad population groups (African Americans, Asian Pacific Islanders, Caucasians and Hispanics). Peptides of greater than 15 amino acids in length consist of adjacent overlapping 15-mers, each with >90% population coverage in the specified broad population groups. Population coverage is further included for all twenty-one detailed population groups.

See supplementary table in Supplementary_S5_Immunogenic_peptides.xlsx.

***Supplementary Table 6: Comparison of HLA diversity among simulated HLA genotype datasets for four broad population groups.***

To explore the impact of HLA genotypic diversity in SARS-CoV-2 proteome immunogenicity, we generated 10 random replicates of HLA genotype datasets encompassing 1,000, 5,000 and 10,000 simulated individuals for each broad population group (African Americans, Asian Pacific Islanders, Caucasians and Hispanics). The number of unique haplotypes required to reach cumulative frequencies of 25%, 50%, 75%, 90% and 95% illustrates that the data follows a power-law distribution, and therefore was compared using the geometric mean to account for significant data skew (using the scipy.stats.mstats.gmean script in Python[6]). This demonstrated comparable results between repeats of the same sample size and population group. Calculation of the alpha parameter value and standard error (sigma) was performed using Powerlaw[7, 8]. This showed that the alpha fit was stable between replicates with a sample size of 10,000, and confirmed that analysis of population samples of size 10,000 was most representative for comparing HLA diversity among populations.

See supplementary table in Supplementary_S6_NMDP_HLA_Class_II_genotypic_diversity_stats.xlsx

***Data availability***

All source data on haplotype frequencies covering 99% of each population group, and all HLA genotype datasets for each population examined, including replicates, have been deposited and are publicly available at Mendeley Data: doi:10.17632/545r9cggzf.1
